# TNF signaling maintains local restriction of bacterial founder populations in intestinal and systemic sites during oral *Yersinia* infection

**DOI:** 10.1101/2025.02.26.639286

**Authors:** Stefan T. Peterson, Katherine G. Dailey, Karthik Hullahalli, Daniel Sorobetea, Rina Matsuda, Jaydeen Sewell, Winslow Yost, Rosemary O’Neill, Suhas Bobba, Nicolai Apenes, Matthew E. Sherman, George I. Balazs, Charles-Antoine Assenmacher, Arin Cox, Matthew Lanza, Sunny Shin, Matthew K. Waldor, Igor E. Brodsky

## Abstract

Enteroinvasive bacterial pathogens are responsible for an enormous worldwide disease burden that critically affects the young and immunocompromised. *Yersinia pseudotuberculosis* is a Gram-negative enteric pathogen, closely related to the plague agent *Y. pestis,* that colonizes intestinal tissues, induces the formation of pyogranulomas along the intestinal tract, and disseminates to systemic organs following oral infection of experimental rodents. Prior studies proposed that systemic tissues were colonized by a pool of intestinal replicating bacteria distinct from populations within Peyer’s patches and mesenteric lymph nodes. Whether bacteria within intestinal pyogranulomas serve as the source for systemic dissemination, and the relationship between bacterial populations within different tissue sites is poorly defined. Moreover, the factors that regulate *Yersinia* colonization and dissemination are not well understood. Here, we demonstrate, using Sequence Tag-based Analysis of Microbial Populations in R (STAMPR), that remarkably small founder populations independently colonize intestinal and systemic tissues. Notably, intestinal pyogranulomas contain clonal populations of bacteria that are restricted and do not spread to other tissues. However, populations of *Yersinia* are shared among systemic organs and the blood, suggesting that systemic dissemination occurs via hematogenous spread. Finally, we demonstrate that TNF signaling is a key contributor to the bottlenecks limiting both tissue colonization and lymphatic dissemination of intestinal bacterial populations. Altogether, this study reveals previously undescribed aspects of infection dynamics of enteric bacterial pathogens.

**Importance:** Bacterial escape from the intestine can lead to severe disease, including sepsis, organ damage, and death. However, the intestinal bacterial population dynamics governing the colonization of mucosal and systemic tissues and the intestinal sites that seed systemic spread are not clear. *Yersinia pseudotuberculosis* is a rodent and human intestinal pathogen closely related to the plague agent and provides a natural rodent-adapted model to study systemic bacterial dissemination. Our findings define the infection dynamics of enteric *Yersinia* and the impact of the innate immune system on *Yersinia* colonization of the intestine and systemic organs.

## Introduction

Enteric bacterial pathogens account for an estimated 3.6 million illnesses per year in the United States and are responsible for 64% of hospitalizations and 64% of deaths caused by contaminated food^1^. The burden of bacterial diarrheal pathogens is even greater in developing regions^2^. Several enteric bacterial pathogens can disseminate from the intestine and replicate within systemic sites, including the liver, spleen, and lungs, due to the presence of virulence factors or infection of immunocompromised individuals ^3,4^. Disease outcomes can be particularly severe in settings of systemic bacterial spread^4^. Despite the importance of systemic dissemination in the lifecycle of enteroinvasive pathogens, the mechanisms leading to extraintestinal spread, the sites of intestinal replication that seed systemic sites, and the host-derived bottlenecks that limit these processes remain incompletely defined.

Population and dissemination dynamics within a host cannot be inferred by quantifying bacterial burdens in tissues alone. Barcoded libraries in which unique nucleotide sequences are inserted into a fitness-neutral site of the bacterial chromosome create allelic diversity in otherwise identical cells, providing a method to dissect the early events that lead to systemic dissemination^5^. This approach allows us to monitor individual clones during infection, thereby enabling quantification of the number of bacteria from the inoculum that give rise to the population at the site of infection, known as the founding population^6^. In addition, analysis of the sequence and abundance of individual clones from the library in different tissue sites by high-throughput sequencing enables robust description and quantification of dissemination dynamics.

*Yersinia* are Gram-negative bacteria that cause disease ranging from plague to severe gastroenteritis^7^. *Y. pseudotuberculosis* (*Y. ptb*) is a natural pathogen of humans and wild animals^8^ that causes self-limiting gastroenteritis and lymphadenitis in immunocompetent patients^9^. However, in immunocompromised individuals, such as those with hemochromatosis or human immunodeficiency virus infection, extraintestinal dissemination and septicemia can occur, causing hepatic or splenic abscesses, septic arthritis, or meningitis^10^. As *Y.ptb* is a natural rodent pathogen, experimental infection of mice provides a tractable model to study mechanisms of systemic dissemination following gastrointestinal inoculation.

While the molecular pathways involved in *Y. ptb*-host cell interactions are well-known^11–13^, little is known about the tissue-level dynamics of the *Y. ptb* populations that establish acute infection. A complex and important series of events influence the establishment of an intestinal infection, including a pathogen’s ability to survive or subvert the low pH of the stomach, endogenous intestinal microbiota, and antimicrobial peptides and other innate immune factors present along the intestinal epithelium^14,15^. These barriers are augmented by immune cells and factors within the lamina propria that influence sites of bacterial infection as well as the trafficking of bacteria within and outside of intestinal tissue. The relative contributions of these bottlenecks to infection dynamics remain largely uncharacterized in many enteric infections, including *Y. ptb*.

Following oral *Yersinia* infection in mice, the Peyer’s patches and mesenteric lymph nodes harbor high bacterial burdens prior to systemic infection^16,17^. Peyer’s patches and mesenteric lymph nodes do not share founding populations with the liver and spleen during *Y. ptb* infection^16^. Instead, the liver and spleen harbor *Y. ptb* populations that closely resemble a pool of *Y. ptb* in the ileal portion of the small intestine. However, the precise anatomic location of these bacteria within the intestine has remained elusive. We recently identified and characterized intestinal microcolonies of *Y. ptb* within pyogranulomatous lesions in the lamina propria of the small intestine^18^. Moreover, we found that the pleiotropic inflammatory cytokine TNF is critical for robust formation of intestinal pyogranulomas, control of *Y. ptb* burden, and host survival^19^.

To address the contribution of intestinal pyogranulomas to either control or dissemination of enteric *Yersinia* and to dissect the relationship between bacterial populations within mucosal intestinal sites and systemic tissues, we employed Sequence Tag-based Analysis of Microbial Populations in R (STAMPR)^6^ together with a newly-generated *Y. ptb* barcoded library containing approximately 56,000 unique barcodes. We find that a surprisingly small population of founder bacteria independently seed sites of the small intestine, and that each pyogranuloma is generated from the expansion of a single bacterium. Notably, Peyer’s patches and pyogranulomas contain high numbers of bacteria, yet are restrictive sites impeding bacterial dissemination, as they do not share bacterial populations with any other tissue sites. Our data indicate that a unique bacterial population likely disperses systemically via hematogenous spread and seeds the systemic infection sites independently of mucosal intestinal populations. Intriguingly, our data further demonstrate that tumor necrosis factor (TNF) signaling restricts both the colonization of tissues by limiting founding population sizes and impedes their dissemination, particularly through the lymphatics. Altogether, this study uncovers key aspects of intestinal bacterial dissemination following oral infection and demonstrates that despite initially supporting bacterial replication, mucosal intestinal tissue sites are highly restrictive to bacterial dissemination.

## Results

### Bacterial tissue burdens are driven by extensive replication of small numbers of initial founders

Previous work identified non-Peyer’s Patch ileal tissue as a source for extraintestinal spread of *Y. ptb* to systemic sites^16^. This, combined with our finding of intestinal pyogranulomas containing *Y. ptb* microcolonies following oral inoculation^18^, raised the possibility that intestinal pyogranulomas might represent the source of systemic bacterial populations. To test the contribution of pyogranulomas in bacterial spread and dissect the factors that affect infection dynamics overall, we generated a library of otherwise isogenic, barcoded *Y. ptb* in the IP2777 strain background. This strain was originally isolated from a septicemic patient at Institute Pasteur^20^ and has been extensively characterized in murine models of oral *Y. ptb* infection^18,19,21^. To generate this library, we inserted random 25-nucleotide sequences downstream of the *glmS* locus using the Tn7 system^22^. Sequencing the library (IP2777-STAMPR) indicated that it contained ∼56,000 unique barcodes, and STAMPR analysis of the library revealed that it could be used to determine founding population sizes of up to 10^6^ founders (Supp. Fig. 1A). Mice infected with 2 x 10^8^ CFU IP2777-STAMPR showed similar burdens in intestinal and systemic tissues compared to our previous studies of mice infected with the parental IP2777 strain^18^ (Fig. 1A), indicating that insertion of the barcodes did not affect the colonization capacity of IP2777. Consistent with our previous findings, we observed that intestinal pyogranulomas (PG+) harbor greater than 100-fold higher burden of *Y. ptb* compared to adjacent non-pyogranulomatous tissue (PG-, Fig. 1B).

**Figure 1.**
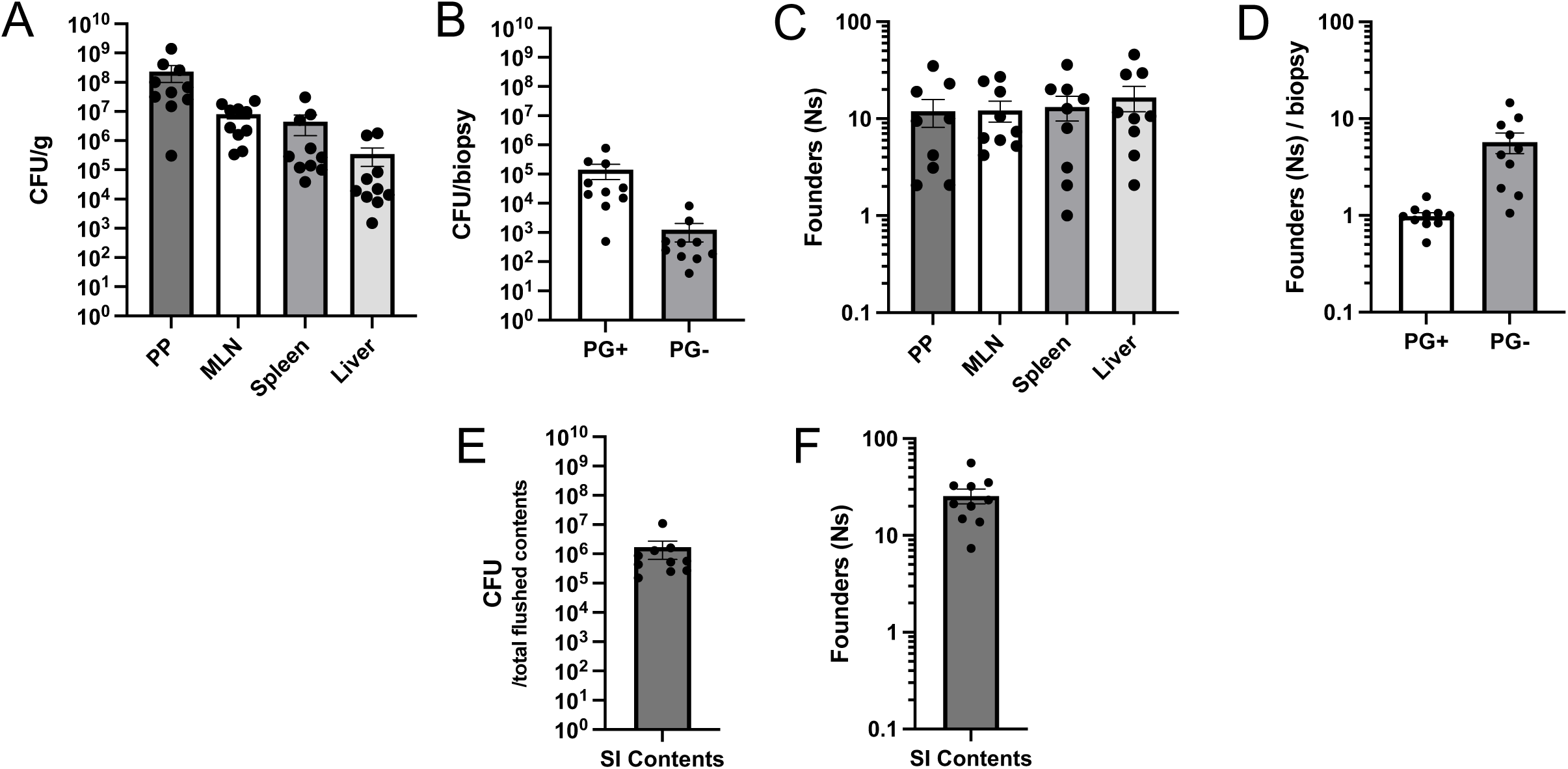
Bacterial tissue burdens are driven by extensive replication of small numbers of initial founders. (A) Bacterial burdens in indicated tissues at day 5 post-infection. (B) Bacterial burdens in small intestinal PG+ and PG- tissue isolated day 5 post-infection. Each circle represents the total CFU of 5-10 pooled punch biopsies divided by number of pooled punch biopsies from one mouse. (C) Founding population (Ns) in indicated tissues at day 5 post-infection. (D) Founding population (Ns) in small intestinal PG+ and PG- tissue isolated day 5 post-infection. Each circle represents the total Ns of 5-10 pooled punch biopsies divided by number of pooled punch biopsies from one mouse. (E) *Y. ptb* bacterial burden in flushed small intestinal (SI) contents at day 5 post-infection. (F) Founding population (Ns) in flushed SI contents at day 5 post-infection. For all data, circles represent one mouse, bars are mean ± standard error of the mean (SEM), and data are pooled from three experiments.

Tissues from mice infected with IP2777-STAMPR were collected, deep sequenced at the barcode locus, and *Y. ptb* populations were analyzed using STAMPR pipeline^6^. Bottlenecks were quantified by calculating the founding population, the number of bacteria from the inoculum that give rise to the observed population at a site of infection, via a metric called Ns. Ns calculation is achieved through a computational resampling-based approach that simulates infection bottlenecks and determines the sampling depth required to observe a given number of barcodes seen in a tissue sample. High Ns values indicate less restrictive infection bottlenecks than lower Ns values. Notably, we observed that the founding populations in both intestinal and systemic sites were very small fractions of the inoculum size: the Peyer’s Patches, mesenteric lymph nodes (MLN), spleen, and liver each had approximately 12-17 founders on average per site, with a range of 1-40 in any given mouse (Fig. 1C). This average represents a sharp narrowing of the population, with only 0.000008% of the clones in the inoculum being found in the tissues following infection. Intestinal biopsies containing pyogranulomatous or non-pyogranulomatous tissues (PG+ and PG-) were pooled per mouse, and the size of founding populations at these sites was determined by dividing the Ns by the number of biopsies taken from each respective mouse (Fig. 1D). In the non-granulomatous intestinal tissue, similar to other tissues, we observed an oligoclonal population with approximately 8-10 unique founders per PG- biopsy. In contrast, pyogranuloma tissue primarily contained a single founder per biopsy, indicating that bacterial populations within intestinal pyogranulomas are clonal. We considered the possibility that a large portion of the inoculum might remain within the intestinal lumen rather than initially colonizing the host tissue. However, the *Y. ptb* burden in the small intestinal (SI) contents during acute infection contained only 10-40 founders (Fig. 1E-F). Surprisingly, the low number of founders did not result from rapid killing of the inoculum nor colonization resistance; high burdens in the stool 1-hour and 6-hours post-inoculation show that a large number of founders survive and traverse the entire gastrointestinal tract (Supp Fig. 1B-C). Furthermore, broad-spectrum antibiotic pretreatment increased bacterial burdens and the number of founders in the intestinal lumen and in non-pyogranulomatous intestinal tissues but had minimal effect on founding populations or burdens at other intestinal and extraintestinal sites, with the exception of the MLN, which had a higher Ns (Supp Fig. 1D-J). Therefore, microbiota shape the bottleneck in the intestinal lumen but not in other compartments. Altogether, these data demonstrate that *Y. ptb* face extremely tight immune bottlenecks for intestinal colonization and tissue infection and that very small populations are responsible for all subsequent replication that occurs in tissues.

### Peyer’s patches and intestinal pyogranulomas contain distinct bacterial populations

*Yersinia* microcolonies in splenic abscesses following intravenous injection of a mixed population of both GFP- and mCherry-expressing bacteria were found to be clonal^17,23^. We observed that the number of unique bacterial founders in pooled intestinal pyogranuloma samples was similar to the total number of biopsies collected, suggesting that bacterial populations within individual pyogranulomas are also clonal. To definitively test whether individual intestinal pyogranulomas contained clonal bacterial populations, we isolated individual pyogranuloma (PG+), Peyer’s patches (PP), and adjacent non-granulomatous tissue (PG-) biopsies from 4 mice and mapped them along the length of the gastrointestinal tract (Supp. Fig. 2A-D). CFU and Ns per biopsy were similar when individual biopsies were harvested (Fig. 2A-B) or when biopsies were pooled and processed together (Fig. 1A-D). Individual PP and PG- typically harbored multiple disparately abundant clones, with 76% and 100% having greater than 1 barcode, respectively (Fig. 2B-C, Supp. Fig. 2E-G). In contrast, individual PG+ primarily only contained a single barcode (58%). These data indicate that intestinal pyogranulomas primarily harbor clonal populations of *Y. ptb*.

**Figure 2.**
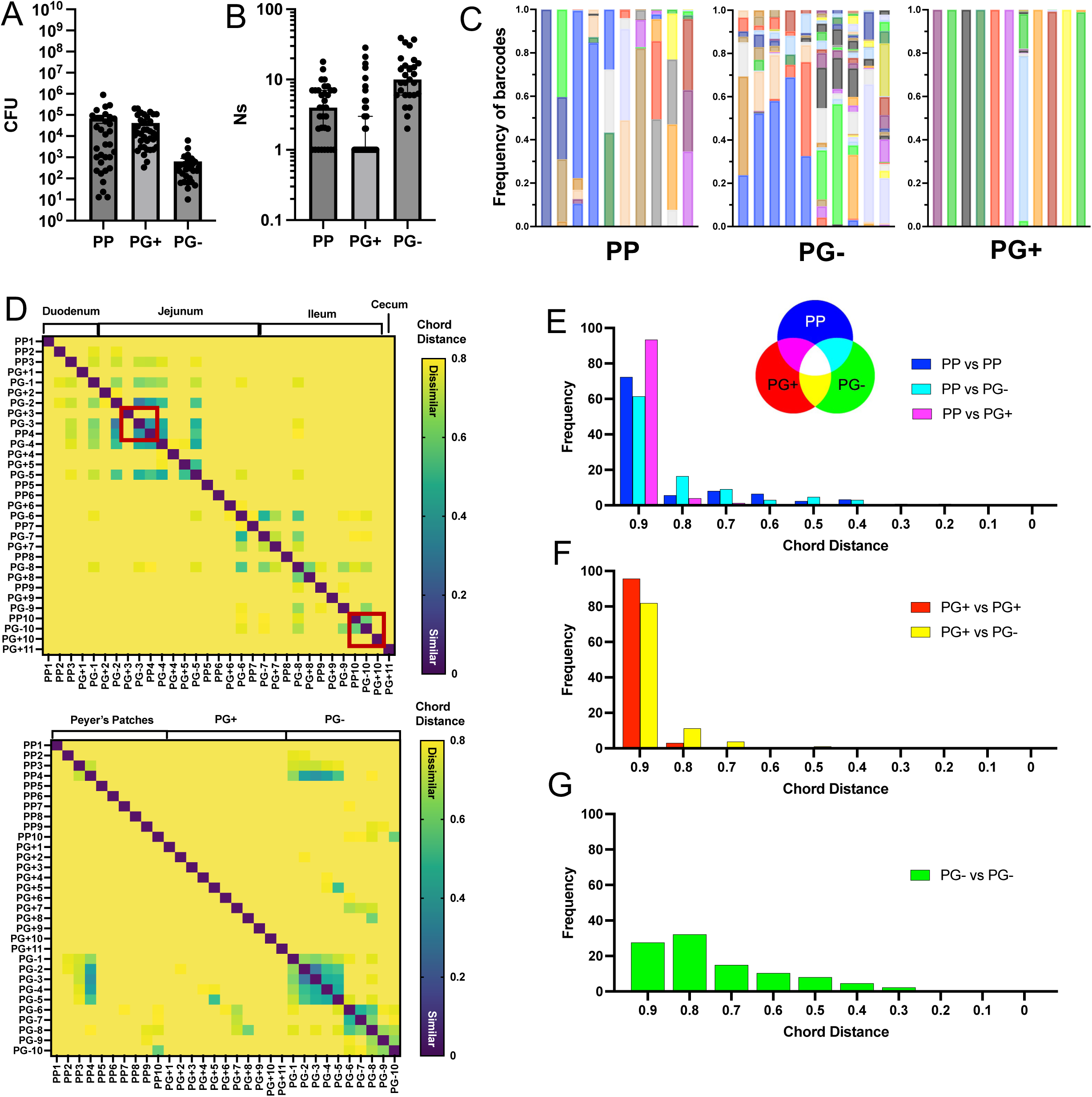
Peyer’s patches and intestinal pyogranulomas contain distinct bacterial populations. (A) Bacterial burdens and (B) founding population (Ns) in small intestinal PP, PG+, and PG- tissue isolated day 5 post-infection. Each circle represents one biopsy. Bars represent (A) mean ± SEM or (B) median ± 95% CI. Pooled data from four mice from one experiment. (C) Frequency of barcodes per biopsy for mouse 1. Each bar represents one biopsy, and each color represents one barcode. (D) Similarity between *Y. ptb* populations in each biopsy as assessed by chord distance (CD) for mouse 1, depicted as (TOP) samples ordered by location along small intestine or (BOTTOM) samples ordered by biopsy type. Red boxes highlight two examples of sharing dynamics where PP share with adjacent PG-, but not with adjacent PG+. (n=4, one experiment, equivalent data of mice 2-4 is depicted in Supp. Fig 2E-J). (E) Histograms showing the frequency of binned CD values for each comparison: (E) PP vs PP (blue), PP vs PG- (cyan), and PP vs PG+(magenta); (F) PG+ vs PG+ (red) and PG+ vs PG- (yellow); and (G) PG- vs PG- (green). For E-G, color-coding of comparisons is diagrammed by triple Venn diagram with specific comparisons highlighted in the figure key, bin width = 0.1, and the center of bins are shown on x-axis. Data is cumulative from 4 mice, one experiment.

To test the possibility that bacteria within pyogranulomas might derive from bacterial populations in Peyer’s patches or adjacent non-granulomatous tissue, or conversely, that bacteria within pyogranulomas might seed bacterial populations in Peyer’s patches, we compared the bacterial populations along the length of the GI tract using the Cavalli-Sforza Chord Distance analysis of the STAMPR pipeline (CD)^6^. This metric takes into account barcode identity and relative abundance in populations being compared. While the highest possible CD between two populations is (2*√2)/II ≈ 0.9, a CD of ∼0.8 or higher indicates that two populations are meaningfully dissimilar. A lower CD indicates populations are more similar to one another, with ∼0.2 being profoundly similar. Largely, we observed that adjacent regions of the intestine were more likely to have lower CD, and thus harbor similar *Y. ptb* populations, than physically distant biopsies (Fig. 2D, Supp. Fig. 2H-J). We did not find a difference in population sharing dynamics based on the intestinal region (duodenum, jejunum, ileum, Supp. Fig. 2K), indicating that intestinal geography did not impact the dynamics of bacterial populations.

*Y. ptb* is thought to translocate into intestinal tissue through M cells that predominantly overlay Peyer’s Patches^24^. If *Y. ptb* initially replicated in Peyer’s Patches and then went on to spread to pyogranulomas and adjacent non-pyogranulomatous tissue, we would expect PP and other biopsies to harbor more similar populations of *Y. ptb*, and, therefore, lower CD. Interestingly, however, PP are mostly dissimilar to other intestinal biopsies (Fig. 2E). However, we did observe small numbers of PP that have similar *Y. ptb* populations (lower CD) to other PP and uninflamed intestinal sites (PG-). In contrast, over 93% of PP to PG+ comparisons are completely dissimilar, as indicated by CD values greater than 0.8 (Fig. 2E). For example, in the mouse depicted in D, we observed shared clones between Peyer’s patches and adjacent uninflamed tissues, but no sharing with adjacent pyogranulomas (Fig. 2D, red boxes). Because the *Y. ptb* populations in intestinal pyogranulomas are clonal and those in Peyer’s patches are oligoclonal, CD could be masking instances where a founder is shared but represents a small proportion of the clones in the Peyer’s patch. Therefore, we enumerated the frequency of instances where a pyogranuloma founder was also present in a Peyer’s patch of the same mouse. Sixty-six percent of pyogranulomas did not have a *Y. ptb* founder that was shared with any Peyer’s patches (Supp. Fig. 2L). Together, these results indicate that *Y. ptb* populations in Peyer’s Patches and intestinal pyogranulomas are generally distinct from one another.

We observed regions of the intestine containing multiple pyogranulomas in proximity (Supp. Fig. 2A-D), raising the question of whether closely apposed pyogranulomas might also be seeded by common founder bacteria. However, there was complete dissimilarity among PG+, as indicated by CD values greater than 0.8 (Fig. 2F), and only two instances where adjacent pyogranuloma biopsies shared *Y.ptb* founders (Supp. Fig. 2H-I). These data indicate that intestinal pyogranulomas are seeded by independent, unique, founder bacteria from within the initial inoculum pool. PG+ *Y. ptb* populations were largely dissimilar to populations in PG- biopsies (Fig. 2F). However, while pyogranulomas rarely share clones with Peyer’s patches or other pyogranulomas, 60% of PG+ share their top barcode with one or more PG- biopsies (Supp. Fig 2L), indicating that the adjacent non-granulomatous tissue could be a source for the bacterial populations within pyogranulomas or that *Y. ptb* in pyogranulomas can escape into the lamina propria. Interestingly, we observed a wide CD distribution for comparisons between individual PG- biopsies, with 30% of biopsy comparisons less than 0.8 (Fig. 2G). This significantly lower distribution of CDs (Supp. Fig. 2M-N) indicates that distinct areas of non-inflamed intestinal tissue share similar *Y. ptb* populations more than Peyer’s patches or pyogranulomas. Together, these data show that the *Y. ptb* populations that seed Peyer’s patches and pyogranulomas are distinct, with unique clonal populations forming in pyogranulomas.

### Bacterial populations in the spleen and liver are distinct from those in the intestine

Prior studies demonstrated that *Y. ptb* disseminates to systemic sites from an intestinal population that is distinct from Peyer’s patches and mesenteric lymph nodes^16^. We, therefore, sought to determine whether the systemic sites might be colonized by *Y. ptb* originating from the intestinal pyogranulomas. However, as for other intestinal sites, the CD between *Y. ptb* populations in the spleen and PG+ was at or above 0.8 (Fig. 3A-B), indicating that pyogranulomas are unlikely to be the source of *Y. ptb* that give rise to the splenic population. Furthermore, while PG+ biopsy populations had a chord distance with splenic populations that was significantly higher than Peyer’s patches or intestinal luminal contents, these areas also had a CD very close to 0.8, indicating that in general these tissues did not share populations with the spleen (Fig. 3B). Furthermore, the splenic population of *Y. ptb* was dissimilar to those in all gastrointestinal organs, including the stomach, cecum, and colon (Supp. Fig. 3A), displaying a general lack of similarity between gastrointestinal and systemic *Y. ptb* populations.

**Figure 3.**
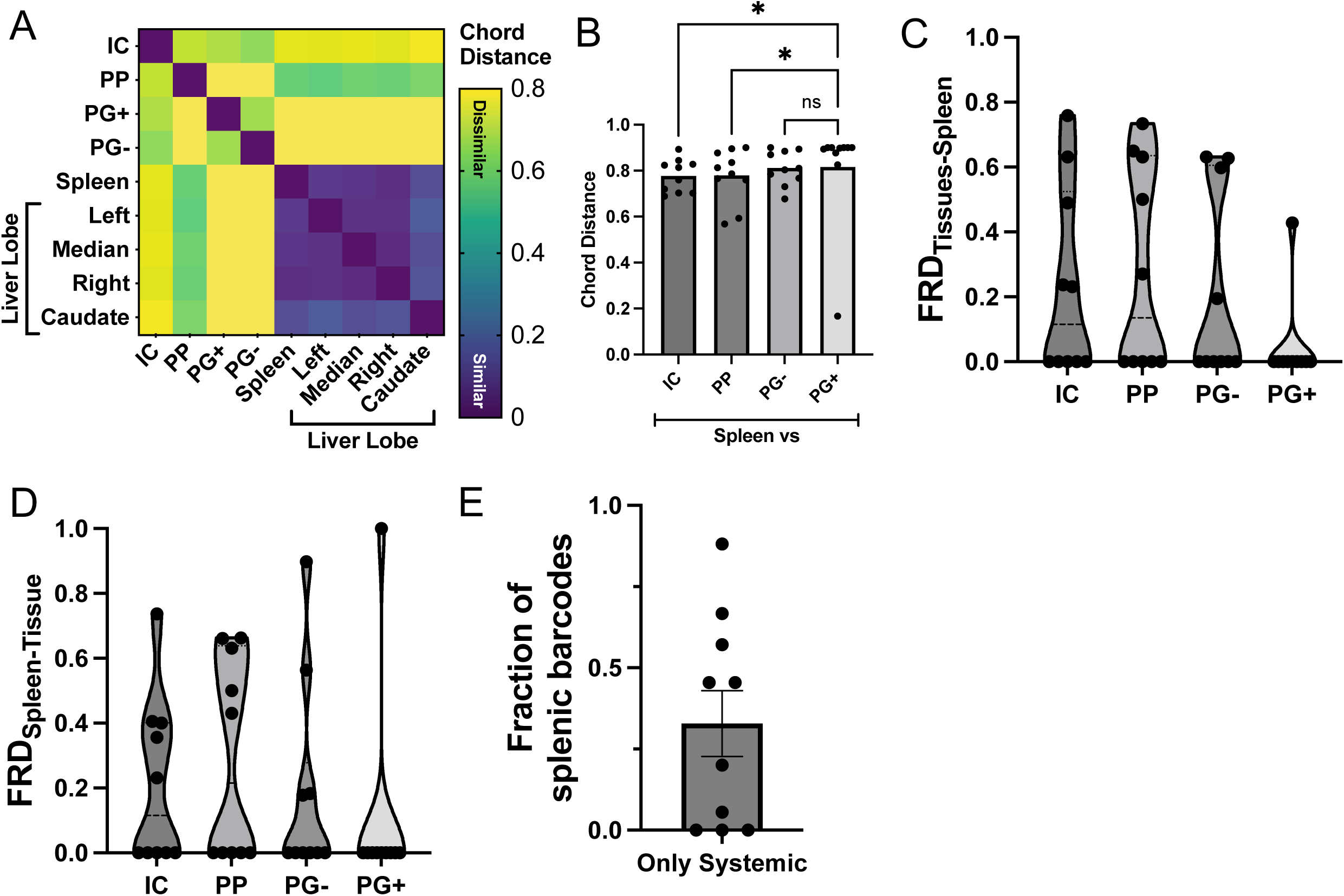
Bacterial populations in the spleen and liver are distinct from those in the intestine. (A) Similarity between *Y. ptb* populations in each sample as assessed by chord distance (CD) for one representative mouse. (B) CD between spleen and indicated samples. (C-D) Prevalence of barcodes from (C) the spleen in other organs (FRD_organ-spleen_) or (D) from various organs in the spleen (FRD_spleen-organ_). (E) Fraction of barcodes from the spleen that are not found in other sites outside of the spleen or liver lobes. All data collected at day 5 post-infection, each circle represents one mouse, n=10, pooled data from three independent experiments, where bars are mean ± SEM and violin plots indicate median ± interquartile range. Statistical significance was calculated using one way ANOVA with post-hoc Dunn’s Multiple Comparison Test, where ns = not significant and * p<0.05.

To test the possibility that individual lobes of the liver might harbor distinct subpopulations, we harvested and processed liver lobes individually and compared them to the spleen and to each other. Notably, the spleen and all liver lobes showed very low CD values relative to one another, indicating that the *Y. ptb* populations in each liver lobe and the spleen are derived from similar clones from the inoculum (Fig. 3A, Supp. Fig. 3B). Additionally, the barcode frequencies in the spleen and liver were similar, with a shared dominant barcode comprising more than 70% of the reads per sample, on average (Supp. Fig. 3C-D). This level of replication by a single founder contrasts with the MLN and intestinal contents, where the most abundant barcode was typically less than half of the barcode reads per sample (Supp. Fig. 3C-D). These data reflect the differential expansion of a single clone across multiple systemic organs, possibly reflecting a “priority effect,” in which early colonization allows a founder replicative advantage over future founders^25^. Collectively, these data indicate that *Y. ptb* populations are readily exchanged between systemic sites and that intestinal sites of *Y. ptb* replication do not serve as sources for dissemination.

We considered the possibility that the CD calculation could be masking populations in different tissues that might have shared founders due to the presence of a single high-abundance founder. Therefore, we used the STAMPR pipeline to calculate a log fraction of the relative numbers of clones shared between two sites, known as FRD^6^. FRD provides insight into the contribution of individual clones to population similarity by quantifying the fraction of barcodes that are shared between two samples. A high FRD_sample1-sample2_ (maximum of 1) indicates that the shared barcodes between sample 1 and sample 2 represent a large fraction of the total barcodes in sample 2. In this example, FRD_sample2-sample1_ could be low, indicating that these same shared barcodes represent only a small fraction of the total barcodes in sample 1.

Notably, in every mouse except one, FRD_PG+-Spleen_ is 0, indicating that none of the barcodes in the spleen are shared with intestinal pyogranulomas (Fig. 3C). Likewise, in most mice, the FRD_Spleen-PG+_ is 0, showing that none of the *Y. ptb* clones in the PG+ biopsies are shared with the spleen. Thus, the intestinal pyogranulomas are not a major source of systemic bacteria. Other intestinal sites have FRDs that are higher than 0, revealing some sharing of barcodes with the systemic sites (Fig. 3C-D). However, no one site consistently shared barcodes with the spleen, suggesting no single site is a common source for dissemination of *Y. ptb* to systemic sites. Lastly, we found that infected spleens can harbor unique *Y. ptb* founders that are not found in any other surveyed tissue (Fig. 3E), suggesting either that *Y. ptb* in systemic organs does not need to replicate in the gastrointestinal tract to disseminate to systemic sites, or that an initial pool from which bacteria might disseminate is no longer present at the time we surveyed the tissue. Together, these data indicate that systemic *Y. ptb* populations do not disseminate from a single consistent intestinal source. Rather, we found that intestinal contents, Peyer’s patches, and non-granulomatous intestinal tissue could occasionally share founders with the spleen; in contrast, we found that intestinal pyogranulomas represent sites of containment that limit spread of bacteria to other intestinal and systemic tissue sites.

### Blood is a conduit for systemic *Y. ptb* populations

Both the lymphatics and bloodstream have been proposed as potential routes for systemic dissemination from the intestine^16,26–28^. We sought to utilize the barcoded *Y. ptb* library to distinguish between these possibilities. Previous studies concluded that *Y. ptb* populations within systemic tissues were distinct from bacterial populations within the MLN^16^. Consistently, we found that the MLN had very high CD to all other sites, including mucosal intestinal sites and systemic tissues (Fig. 4A). Collectively, our findings indicate that sites of bacterial infection and replication in the intestinal mucosa do not serve as direct sources for systemic dissemination and are instead acting as restrictive sites that limit bacterial dissemination to other tissues. Given the lack of similarity between MLN and systemic sites, we also conclude that the lymphatics themselves are unlikely to serve as a conduit for dissemination.

**Figure 4.**
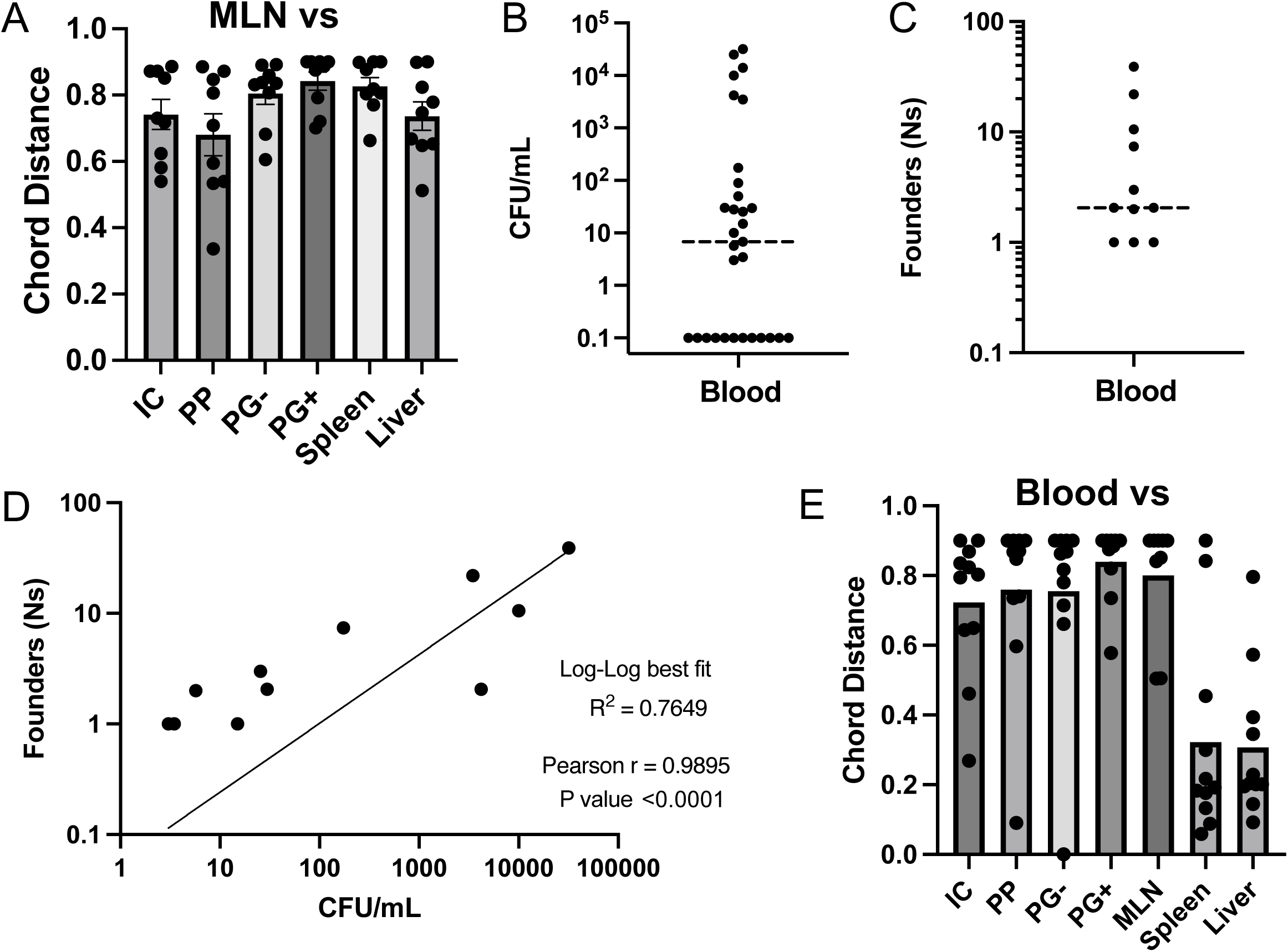
Blood is a conduit for systemic *Y. ptb* populations. (A) CD between the MLN and indicated samples. (B) Bacterial burdens and (C) Founding population (Ns) in blood; blood samples with no CFU are represented as 0.1 CFU. (D) Log-log regression with Pearson’s correlation of fit between Ns and CFU measured in blood. (C-D) Each circle represents one mouse with detectible burden of *Y. ptb* in blood. (E) CD between the blood and indicated samples. Samples labeled liver are showing CD comparisons to the median lobe as representative of all liver lobes. All data were collected at day 5 post-infection, from three-four independent experiments (except B, which has pooled data from seven experiments), where each circle represent one mouse unless indicated otherwise and bars are mean ± SEM.

We therefore considered the possibility that spread of bacteria from the initial founder pool could occur close to the time of initial inoculation via the bloodstream. Interestingly, relatively high numbers of animals (nearly 40%) did not have detectable bacterial levels in the bloodstream, while others contained moderate (10^2^) or high (10^4^) CFU/mL of blood (Fig. 4B). Interestingly, the mice that had detectable bacteria in the bloodstream exhibited a range of unique founders, from a single bacterium in several infected mice to 39 unique founders in one mouse, that showed significant positive correlation with total CFU in blood (Fig. 4C-D). Notably, a comparison of populations in the blood and spleen or liver revealed very low CD between these sites (Fig. 4E), indicating that bacterial populations are shared between the spleen, liver, and bloodstream, consistent with the possibility that dissemination to systemic sites occurs via the bloodstream.

### TNFR1 contributes to *Y. ptb* infection bottleneck and MLN colonization

While it is well appreciated that the immune system plays a key role in controlling pathogen burden, how specific immune components do so is less defined. The immune system might limit the number of founders in a tissue, limit replication of those founders, prevent dissemination from an initial colonization site, or some combination of these activities. TNF is critical for defense against numerous pathogens, including *Yersinia*, where it plays an important role in limiting bacterial tissue burdens and promoting host survival^11,19,29–31^. Consistently, intestinal pyogranulomas in *Tnfr1*^-/-^ mice have elevated bacterial burden and exhibit an increase in tissue necrosis^19^, raising the hypothesis that TNF signaling in pyogranulomas may limit bacterial spread to systemic tissues.

To dissect the impact of TNF signaling on dynamics of bacterial infection as well as the possibility that TNF signaling might be important to contain *Y. ptb* within pyogranulomas, we infected mice with STAMP-IP2777 and quantified founding population sizes and dissemination patterns at day 5 post-infection. While the loss of TNF signaling resulted in no significant effect on luminal *Y. ptb* populations in the small intestine (Supp. Fig. 4A-C), we observed an increase in the number founders as well as an increase in CFU in PP and PG+ tissues (Fig. 5A-B). Notably, intestinal pyogranulomas in *Tnfr1^-/-^* mice harbored oligoclonal *Y. ptb* microcolonies more often than pyogranulomas in wildtype mice (WT), including several mice with 25-50 individual founders. Similarly, PPs of *Tnfr1^-/-^* mice harbored a wider range of founders that WT mice, including several mice with over 100 unique individual founders. These data together indicate that TNF signaling limits initial colonization of intestinal sites, although the variance in founder populations in tissues of individual mice limited the statistical significance of this analysis. Importantly, *Tnfr1^-/-^* mice also had significantly elevated levels of *Y. ptb* in their blood, with all mice carrying at least some bacterial burden in circulation (Fig. 5C). Moreover, we observed a nearly 10-fold increase in the number of unique founders in the blood of *Tnfr1^-/-^* mice (Fig. 5D), indicating that TNF signaling limits colonization of the bloodstream. Intriguingly, in the absence of TNFR1 signaling, the liver and spleen also had higher numbers of unique founders and higher bacterial burdens, indicating that TNF prevents colonization of systemic tissues (Fig. 5E-F). Notably however, TNF signaling does not affect the degree of bacterial replication, as WT and *Tnfr1^-/-^* mice had similar numbers of CFU/founder (Supp. Fig. 4D-F). Together, these data indicate that TNFR1 plays a role in the bottleneck that limits initial colonization of tissue sites.

**Fig. 5.**
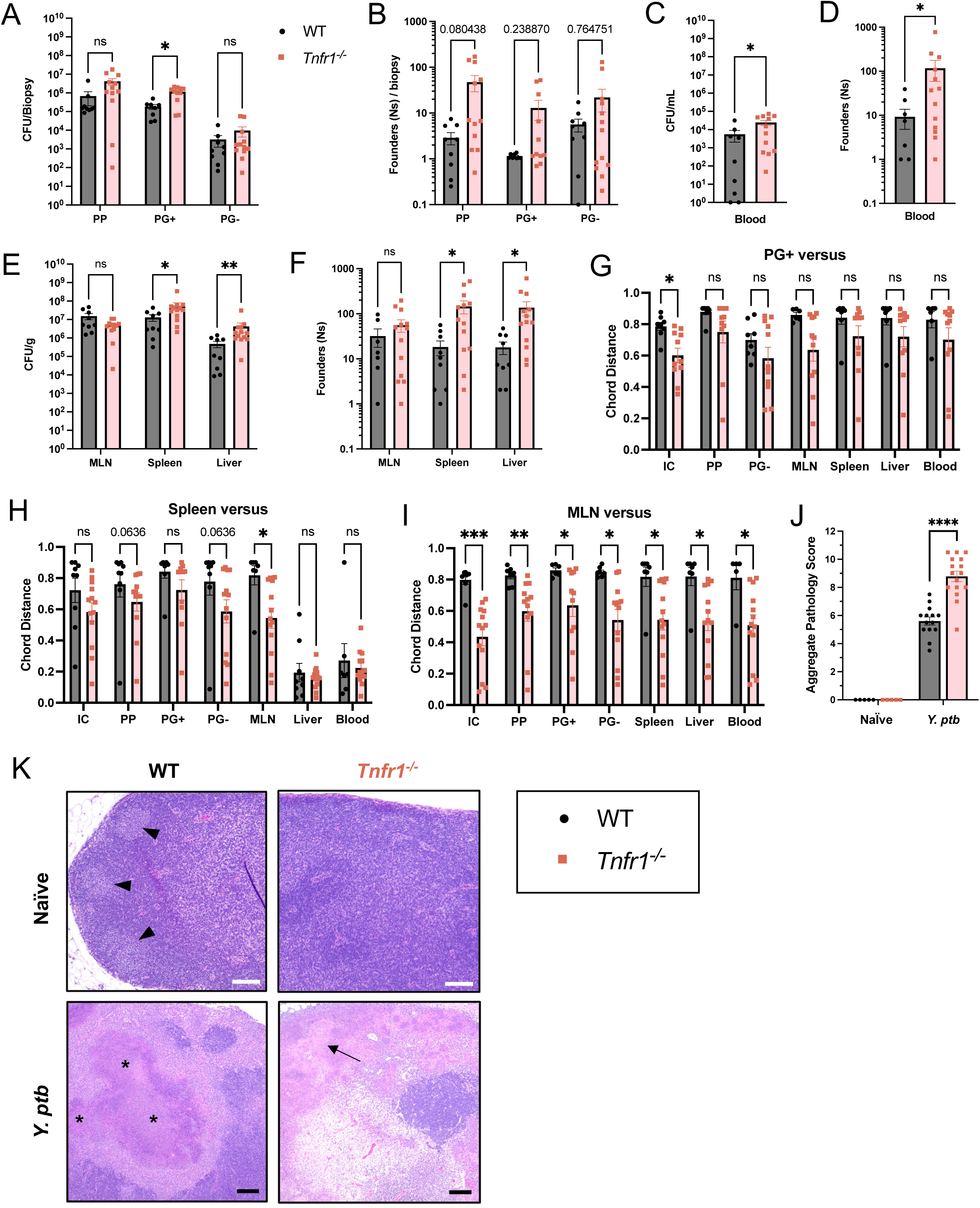
TNFR1 contributes to *Y. ptb* infection bottleneck and MLN colonization. (A) Bacterial burdens and (B) founding population (Ns) in small intestinal PP, PG+, and PG- biopsies at day 5 post-infection, where each symbol represents the total of 5-10 pooled punch biopsies divided by the number of biopsies from one mouse. (C) Bacterial burdens and (D) founding population (Ns) in whole blood. (E) Bacterial burdens and (F) founding populations (Ns) in systemic tissues: mesenteric lymph nodes (MLN), spleen, and liver (median lobe). (G-I) Chord Distance (CD) between (G) PG+, (H) spleen, or (I) MLN and the indicated systemic (MLN, spleen, median liver lobe, blood) and intestinal (PP and PG-) tissues. All data collected on day 5 post-infection and are pooled from three independent experiments. (J-K) H&E-stained paraffin-embedded mesenteric lymph node sections from naïve and *Y.ptb*-infected WT and *Tnfr1^−/−^* mice at day 5 after infection were used for (J) histopathological scoring and (K) imaging. (J) Each mouse was scored between 0 and 4 (minimal to extensive) for the metrics shown in Supp. Fig. 4E-G. Scores for each mouse were added together to obtain the aggregate score shown. (K) Naïve mesenteric lymph nodes from WT mice (LEFT), but not *Tnfr1^−/−^* mice (RIGHT), have lymphoid follicles in the cortex (marked by arrowheads). White scale bars = 100 μm. Infected mesenteric lymph nodes (BOTTOM) show pyogranulomatous lesions in WT mice (marked by asterisks) and necrosis in *Tnfr1^−/−^* mice (marked by arrows). Black scale bars = 200 μm. Histological data are from two independent experiments with representative images shown. Each symbol represents one mouse unless otherwise indicated, bars are mean ± SEM. Statistical significance was calculated using multiple Mann-Whitney tests and ns = not significant, * p<0.05, ** p<0.01, *** p<0.001.

We next examined if the expansion of the colonization bottleneck by TNF signaling deficiency would result in enhanced dissemination either between intestinal sites or from intestinal sites to systemic tissues. Although there was a trend to lower chord distance (increased similarity) between *Y. ptb* populations in intestinal pyogranulomas and other intestinal tissues in *Tnfr1^-/-^* relative to WT mice, this reached significance only for the intestinal luminal contents themselves, suggesting that TNFR1 signaling limits initial colonization of PGs from the intestinal lumen (Fig. 5G). Furthermore, *Tnfr1^-/-^* intestinal PP and PG- populations showed a trend of greater similarity to the spleen populations, with lower CDs than the same comparison in WT mice (Fig. 5H). Critically, the MLN in *Tnfr1^-/-^*mice harbored *Y. ptb* populations with significantly elevated sharing with both intestinal and systemic populations, indicating that TNF signaling limits dissemination between MLN and other tissues (Fig. 5I). Together, these data show that TNFR1 functions as a critical immune factor that imposes infection bottlenecks and controls bacterial dissemination following *Y. ptb* oral infection. Consistent with our previous observation that the architecture of intestinal pyogranulomas is disrupted in the absence of TNF signaling, *Tnfr1^-/-^* mice exhibited disrupted MLN architecture, and increased MLN pathology relative to their wild-type counterparts following *Y. ptb* infection (Fig. 5J-K, Supp. Fig. 4G-I). Intriguingly, MLN architecture is disrupted even in naïve TNFR1-deficient mice, consistent with prior reports that TNF promotes development of B cell follicles and other aspects of lymphoid architecture^32^ (Fig. 5K).

Together, the distinct influences of TNF signaling on intestinal and systemic sites suggests that the role of TNF signaling functions to restrict invasion by individual founders. Thus, the increase in CFU in the liver and spleen may be attributable to the increase in the number of clones that colonize systemic sites. Notably, the increase in burden in systemic tissues in the absence of TNF signaling corresponds with an increase in sharing of *Y. ptb* founders between the systemic sites and the MLN, marking an intriguing role for TNF in lymphatic architecture and bacterial dissemination.

## Discussion

*Y. ptb* is a natural pathogen of both humans and rodents^8^ and has been used extensively to dissect molecular mechanisms of bacterial infection and host immune defense^11–13^. Previous studies indicated that systemic bacterial populations derive from a replicating intestinal pool and are distinct from bacterial populations in Peyer’s patches and mesenteric lymph nodes^16^. Our recent finding that intestinal pyogranulomas that form acutely during *Y. ptb* infection contain large numbers of bacteria^18^ raised the possibility that these pyogranulomas might serve as a source for such systemic populations. To test this possibility, and to further dissect dissemination dynamics of oral *Yersinia* infection, we generated a barcoded population of otherwise isogenic *Y. ptb* and employed the STAMPR analytic pipeline^6^.

The vast majority of the clones in the inoculum initially survive the acidic environment of the stomach and rapidly pass through the intestinal tract within the first 12 hours. Critically, most *Y. ptb* clones fail to colonize any tissues, as only 10-20 founders, on average, colonize the intestinal and systemic sites. This reflects a nearly 10^7^-fold narrowing of the diversity of the initial population at each tissue site, indicating a very tight bottleneck to colonization via the oral route. Notably, each tissue site largely had its own unique subpopulation of bacteria, indicating that most tissue sites are stochastically colonized by independent founder populations. Surprisingly, while antibiotic depletion of the endogenous microbiota in the gut led to an overall increase in *Y. ptb* levels within the intestinal lumen, it did not broadly affect the bacterial tissue burdens, the numbers of unique founders within either intestinal or systemic tissue sites, or spread of bacterial populations between sites, suggesting that the microbiota itself is not a major contributor to the colonization bottleneck for this infection.

Consistent with previous observations^16^, we found that the mesenteric lymph nodes and other intestinal sites did not share founder populations with the systemic populations in the liver and spleen. Moreover, individual pyogranulomas contained largely clonal bacterial populations, in contrast to the other intestinal sites, which contained more oligoclonal populations derived from 2-10 unique bacterial founders. We observed minimal sharing of intestinal pyogranuloma and Peyer’s patch populations, suggesting that Peyer’s patches are not the source for pyogranuloma founders or vice versa. However, we did observe some sharing of Peyer’s patch and pyogranuloma populations with adjacent non-granulomatous intestinal sites. Therefore, while we cannot infer directionality, these findings suggest that there is some limited exchange of bacteria between inflammatory intestinal sites and the adjacent lamina propria. Interestingly, the mesenteric lymph nodes contained an entirely unique population of bacteria that was not shared with any other mucosal intestinal tissue, indicating that in wildtype mice, the MLN is a restrictive site that serves as a dead-end for these bacteria. Notably, both LTβ-deficient mice, which lack Peyer’s patches, and LTα-deficient mice, which lack both Peyer’s patches and MLN, still have systemic *Yersinia* burdens following oral infection^16,33^, further supporting the model that intestinal *Yersinia* disseminate to systemic tissues via an alternate route.

While we were unable to detect the sharing of bacterial populations between intestinal and systemic sites, the liver and spleen populations had significant similarity with one another, indicating either that bacterial founders initially colonize one of these sites and subsequently disseminate to the other or that different subsets of the initial pool independently colonize them and subsequently freely disseminate between the two organs. We also observed that the liver and splenic populations exhibited significant sharing with the bacterial population in the bloodstream. Importantly, while overall bacterial numbers and unique founders in the blood were low to undetectable in most mice, those in which we could isolate bacteria from the bloodstream exhibited significant sharing with systemic sites. Interestingly, bloodstream CFU data could be divided into three subsets of animals: mice with undetectable bacterial burdens in the bloodstream, mice with low-intermediate levels (10-100 CFU/mL of blood), and mice with much higher burdens averaging 10^4^ CFU/mL. Regression analysis revealed that bacterial numbers in the blood strongly correlated with the number of unique founders in the blood suggesting that blood burden is determined by entry of unique founders into the circulation. Hematogenous spread of gastrointestinal pathogens has been reported in *Salmonella* infection^34,35^, suggesting a common route of transport to the systemic sites for both intracellular and extracellular pathogens. Ultimately, these findings, alongside recent work with *Klebsiella pneumoniae*^36^ and *Salmonella enterica*^35^, suggest that systemic dissemination doesn’t require replication in the upstream site, which is important for our understanding how bacterial infections progress.

While burdens of *Y. ptb* in systemic circulation were often below the level of detection in WT mice, in mice lacking TNFR1, *Y. ptb* was robustly detected in circulation with significantly higher burdens in systemic organs. These higher burdens correspond to higher numbers of founders in systemic sites, Peyer’s patches, and intestinal pyogranulomas, indicating that TNF signaling controls *Y. ptb* capacity to establish founder populations across tissues. Surprisingly, dysregulated pyogranulomas in *Tnfr1^-/-^* mice can harbor oligoclonal *Y. ptb* microcolonies. However, *Y. ptb* populations in *Tnfr1^-/-^* pyogranulomas were not significantly more similar to other infected tissue sites than wildtype pyogranulomas, suggesting that even in TNFR1-deficient animals, intestinal pyogranulomas restrict dissemination of the bacteria. Notably, in the absence of TNFR1, Peyer’s patches and non-inflamed tissues of the intestine showed a trend to increased sharing with systemic sites. However, the largest impact of TNF signaling deficiency on *Y. ptb* intra-host population dynamics was in the MLN, which consistently showed significantly higher levels of sharing with every other tissue of *Y. ptb* colonization across multiple animals. These findings suggest that TNF signaling in MLNs restricts systemic dissemination of intestinal bacterial pathogens. Future studies will investigate the mechanism of TNF’s effect on lymphatic spread of *Y. ptb* and the potential role of IL-1 downstream of TNF signaling^19^, in infection bottlenecks during intestinal *Y. ptb* infection.

Overall, our results support a model (Fig. 6) where much of an initial infecting inoculum passes through the stomach and arrives in the intestines shortly after inoculation. The pyogranulomas that form in the small intestine during acute infection are seeded by unique single founders that establish localized replicative niches. In areas of the intestine lacking pyogranulomas, *Y. ptb* clones replicate to a lesser extent but can be found at multiple sites across the intestinal tract. Our findings also imply that on the order of 10-20 initial founders translocate across the intestinal barrier and reach systemic circulation, which mediates their trafficking to and between the liver and spleen. Our evidence suggests that the initial barriers to colonization of tissues are partially controlled by TNF signaling, which also limits translocation to intestinal tissue and systemic sites, particularly from the MLN. Our study reveals key aspects of *Yersinia* population dynamics within the host and highlights the key role of the innate immune system in controlling infection by limiting extraintestinal dissemination patterns.

**Figure 6.**
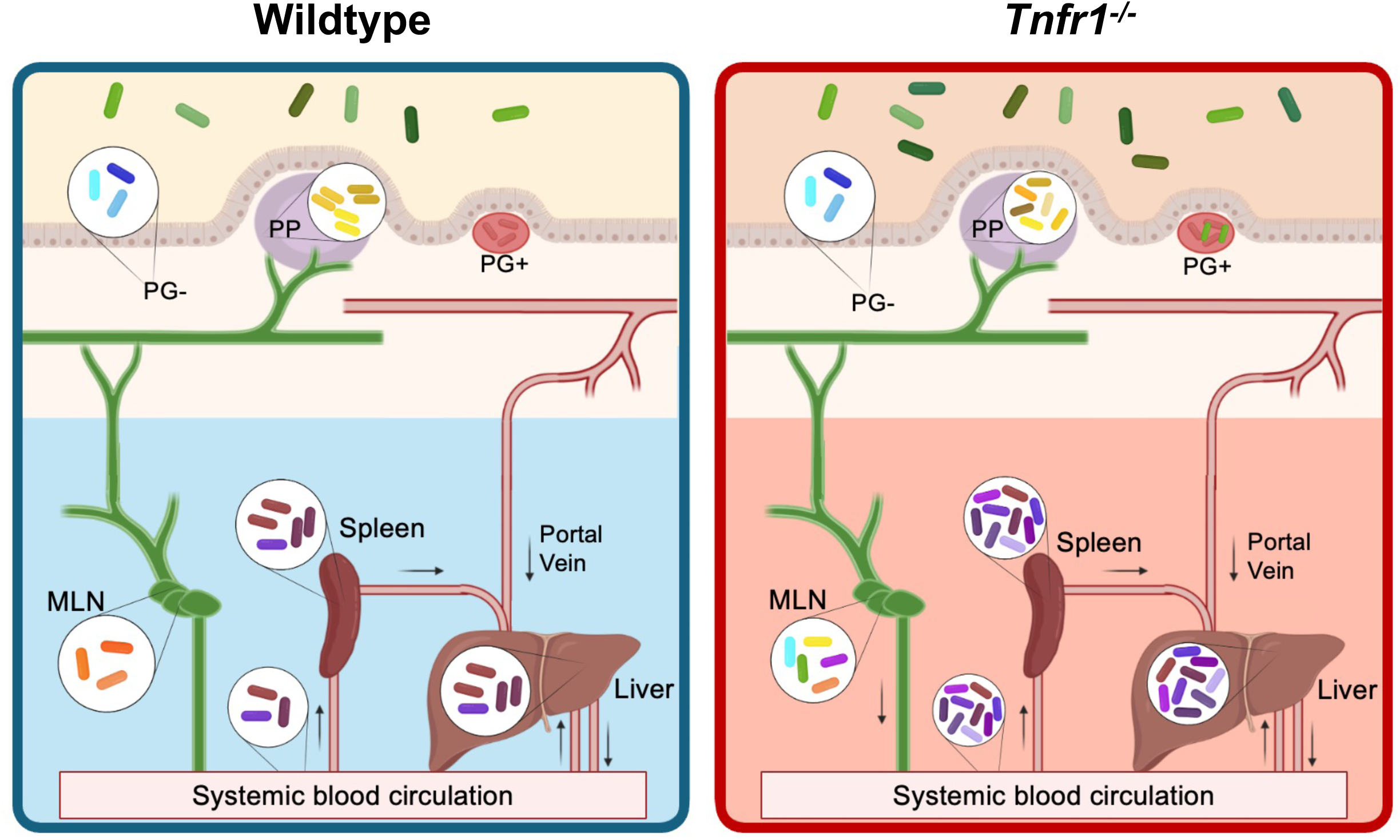
Barcoded *Y. ptb* reveal unique subpopulations across organs, hematogenous spread, and role for TNFR1 in lymphatic restriction of clones. (LEFT) Wildtype mice are able to control bacterial burdens by maintaining tight bottlenecks to infection. A small number of clones (each depicted by a unique color) are able to seed infection by invading intestinal tissue and replicating or disseminating directly systemic organs via the blood. We found that intestinal pyogranulomas (PG+) harbor a clonal population of *Y. ptb* that is contained, thus not the source for systemic dissemination. Meanwhile, mesenteric lymph nodes (MLN) harbor a small population of *Y. ptb* clones that is restricted to this site. (RIGHT) *Tnfr1^-/-^* mice are not able to control bacterial burdens because of wider bottleneck allowing more clones to colonize sites of infection. Necrotic PG+ biopsies occasionally harbor oligoclonal populations of *Y. ptb*. MLN samples reveal founders that are shared with other sites of infection.

## Methods

### Mice

C57BL/6 (Strain #000664) wildtype were acquired from the Jackson Laboratory. *Tnfr1^−/−^*mice were previously described^19,30^. All mice were bred at the University of Pennsylvania by homozygous mating and housed separately by genotype. Mice of either sex between 8–12 weeks of age were used for all experiments. All animal studies were performed in strict accordance with University of Pennsylvania Institutional Animal Care and Use Committee-approved protocols (protocol #804523).

### Generation of STAMP library

Wildtype *Y. ptb* (clinical isolate strain 32777 aka IP2777, serogroup O1)^20^ was provided by Dr. James Bliska (Dartmouth College) and previously described^37^. A revTet constitutive mCherry construct (citation) provided by Dr. Kimberly Davis (Johns Hopkins University) was cloned into a previously described neutral site, YPK_2061^38^, using two-step allelic recombination with a plasmid provided by Dr. Joan Mecsas (Tufts University). This IP2777-mCherry was used to create the barcoded library used throughout this manuscript. The growth of this mutant in 2xYT broth was indistinguishable from the wildtype strain from which it was derived by growth curves.

The library “STAMP-IP2777” was created, as described previously^39,40^, using the plasmid donor library pSM1^41^. The pSM1 donor library is composed of ∼70,000 unique plasmids transformed into the donor strain MFDλpir. Each pSM1 plasmid carries a site-specific Tn7 transposon containing a random ∼25 nucleotide barcode adjacent to a kanamycin resistance cassette. The Tn7 transposon system integrates at a neutral site in the genome downstream of the gene *glmS*^22^. Conjugation was used to introduce the pSM1 library into IP2777-mCherry, and transconjugants containing the transposon were selected using triclosan and kanamycin. Transconjugant colonies were pooled in PBS with 25% glycerol and frozen at −80 °C in aliquots to create the library STAMP-IP2777. Sequencing indicated that the library contains ∼56,000 unique barcodes.

### Mouse infections

STAMP-IP2777 was prepared for oral inoculation by resuspending frozen aliquots in liquid 2xYT broth supplemented with 2 μg/mL triclosan (Millipore Sigma) and 100 μg/mL kanamycin (GoldBio). Bacteria was cultured to stationary phase at 28°C and 250 rpm shaking for 16 hours. Following growth, bacteria were pelleted and resuspended in phosphate buffered saline (PBS) at a concentration of 2×10^9^ colony-forming units (CFU) per mL. Mice were fasted for 16 hours and subsequently inoculated by oral gavage with 100 μL of inoculum, therefore 2×10^8^ CFU per mouse. Following inoculation, the dose was determined by serial dilution and plating.

### Sample harvesting

Blood was harvested by cardiac puncture upon euthanasia and collected in 250 U/ml Heparin solution (Millipore Sigma). Small intestines were excised, and luminal contents were flushed with sterile PBS into a 50 mL conical for collection. Intestinal contents were then spun down at 3000 relative centrifugal force (RCF) at 4°C for 10 minutes to pellet bacteria and debris. Supernatant was discarded, and the pellet was resuspended in 1 mL of cold, sterile PBS. Resuspended pellets were then homogenized for 40 seconds with 6.35 mm ceramic spheres (MP Biomedical) using a FastPrep-24 bead beater (MP Biomedical). After flushing, small intestines were opened longitudinally along the mesenteric side and placed luminal side down on cutting boards (Epicurean). Small intestinal tissue containing macroscopically visible pyogranulomas (PG+), adjacent non-granulomatous areas (PG−), and Peyer’s Patches (PP) were excised using a 2 mm-ø dermal punch-biopsy tool (Keyes). Biopsies within each mouse were pooled groupwise, unless otherwise stated. Tissues were collected in 1 mL of sterile PBS, weighed, and homogenized for 40 seconds, as previously described^18,19^.

All samples had 100uL serially diluted tenfold in PBS, plated on LB agar supplemented with 2 μg/ml triclosan and 100 μg/mL kanamycin, and incubated for two days at room temperature. Dilutions of each sample were plated in triplicate and expressed as the mean CFU per gram or per biopsy. Remaining 900uL of homogenized samples were plated on 15 cm dishes with LB agar supplemented with 2 μg/ml triclosan and 100 μg/mL kanamycin and incubated for two days at room temperature.

### STAMP sample processing

*Y. ptb* colonies were washed off plates, collected in PBS with 25% glycerol, diluted in water, and boiled for 15 minutes at 95°C. The barcode-containing region was amplified from the genome using custom forward and reverse primers^35,36,39^. Sequence tag-based analysis of microbial populations (STAMP) was performed as previously described^36^. Sequencing data and original script have been deposited on Dryad and will be publicly available at the date of publication. Any additional information required to reanalyze the data reported in this paper is available from the lead contact upon request.

### Antibiotics treatment

Antibiotics-treated mice were given filter-sterilized water *ad libitum* containing 0.5 g/L Vancomycin (Sigma-Aldrich), 0.5 g/L Neomycin (Sigma-Aldrich), 0.5 g/L Ampicillin (Sigma-Aldrich), 0.25 g/L Metronidazole ((Sigma-Aldrich) for 7 days. Antibiotics were removed from the drinking water for 24 hours prior to inoculation with STAMP-IP2777.

### Histology

Tissues were fixed in 10% neutral-buffered formalin (Thermo Fisher Scientific) and stored at 4°C until further processed. Tissue pieces were embedded in paraffin, sectioned by standard histological techniques, and stained with hematoxylin and eosin (H&E) for subsequent assessment of lymph node architecture and histopathological disease scoring by blinded board-certified pathologists. Tissue sections were given a score from 0 to 4 (healthy to severe) for multiple parameters, including degree of inflammation and degree of free bacterial colonies. Healthy mice were characterized by and subsequently scored as having none or low levels of the parameters described, whereas severely afflicted mice presented with high amounts of the respective parameters.

### Statistics

Statistical analyses were performed using Prism v9.0 (GraphPad Software). Independent groups were compared by Mann-Whitney U test or Kruskal-Wallis test with Dunn’s multiple comparisons test. Statistical significance is denoted as * (p<0.05), ** (p<0.01), *** (p<0.001), **** (p<0.0001), or ns (not significant).

## Acknowledgments

We thank Enrico Radaelli for constructive input and the staff at the PennVet Comparative Pathology Core for their help in preparing and analyzing the histological samples. We thank members of the Brodsky and Shin laboratories for helpful scientific discussions, Dr. Heather L. Rossi for editorial suggestions when drafting the manuscript, and Reyna Garcia Sillas for generation of schematic of intestinal position of biopsies and corresponding clones in Sup Fig 2A-D. This work is supported by NIH/NIAID grants, R01AI128530, R01AI1139102A1, and R01DK123528 (I.E.B.); a Burroughs Wellcome Fund Investigator in the Pathogenesis of Infectious Disease award (I.E.B.); Mark Foundation grant 19-011MIA (I.E.B.); a National Science Foundation Graduate Research Fellowships Program Award (S.T.P.). The veterinary pathologists performing the histopathological analysis are supported by the Abramson Cancer Center Support Grant (P30 CA016520). The scanner used for whole-slide imaging and the image visualization software was supported by an NIH Shared Instrumentation Grant (S10 OD023465-01A1).

## Author contributions

S.T.P. conceived the study, devised and performed experiments, and analyzed data. S.T.P., K.G.D., K.H., D.S., R.M., J.S., W.Y., R.O., S.B., N.A., M.S., and G.I.B. performed experiments. C.-A.A., A.C., and M.L. performed the histology and histopathological scoring. S.S., M.K.W., and I.E.B. acquired funding and provided supervision. S.T.P and I.E.B. wrote the original draft. All authors reviewed and edited the manuscript.

## Supplemental Figure Legends

**Supplemental Figure 1.**
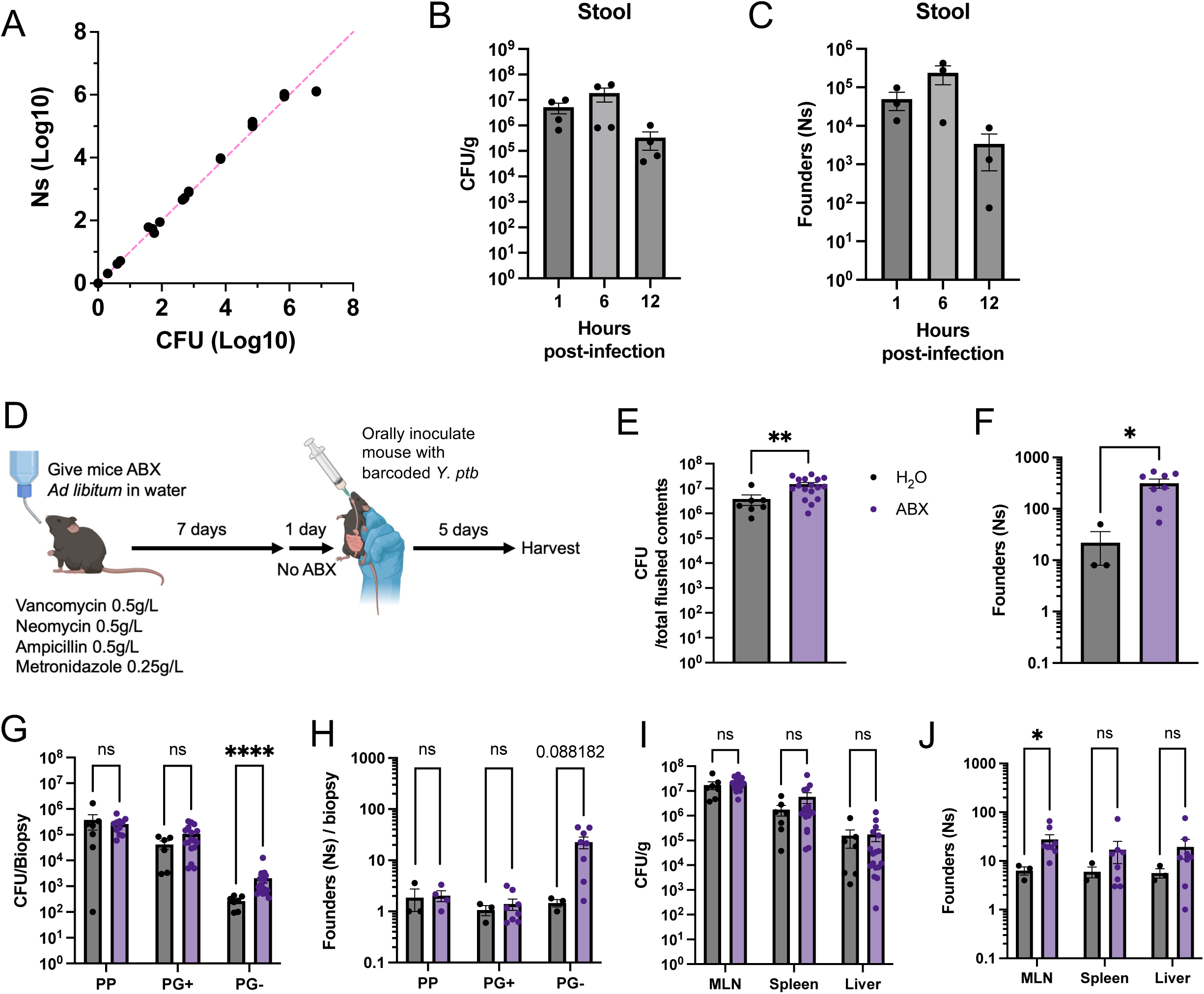
Validation of the *Y. ptb* barcode library and impact of antibiotic pre-treatment on barcoded *Y.ptb* dissemination. (A) The ability to determine the size of the founding population from STAMP libraries was validated in culture by comparing the number of plated colonies (CFU; known number of founders) to the size of the founding population following STAMPR analysis of those plated populations (Ns) for 3 cultures across 9 doses. (B) Bacterial burdens and (C) founding population (Ns) in stool isolated 1-, 6-, or 12-hours post-infection. Pooled data from four mice from one experiment. (D) Graphical representation of the experimental design for antibiotic (ABX) pretreatment: C57BL/6J mice were given either a cocktail of ABX or control sterile lab drinking water ad libitum for 7 days and returned to normal drinking water one day prior to infection. At day 0, mice were inoculated with 2×10^8^ CFU *Y. ptb* library via oral gavage. Mice were euthanized 5 days following the inoculation and *Y. ptb* populations were assessed. (E) Bacterial burden and (F) founding population (Ns) in flushed small intestinal contents. (G) Bacterial burdens and (H) founding population (Ns) in small intestinal PP, PG+, and PG- biopsies. Each circle represents the total CFU of 5- 10 pooled punch biopsies divided by number of pooled punch biopsies from one mouse. (I) Bacterial burden and (J) founding population (Ns) in MLN, spleen, and liver. Unless otherwise indicated, each circle represents one mouse, and all graphs are pooled data from three independent experiments. For bar graphs, bars represent mean ± SEM, and statistical significance was determined using Mann-Whitney tests: ns = not significant, * p<0.05, ** p<0.01, **** p<0.0001.

**Supplemental Figure 2.**
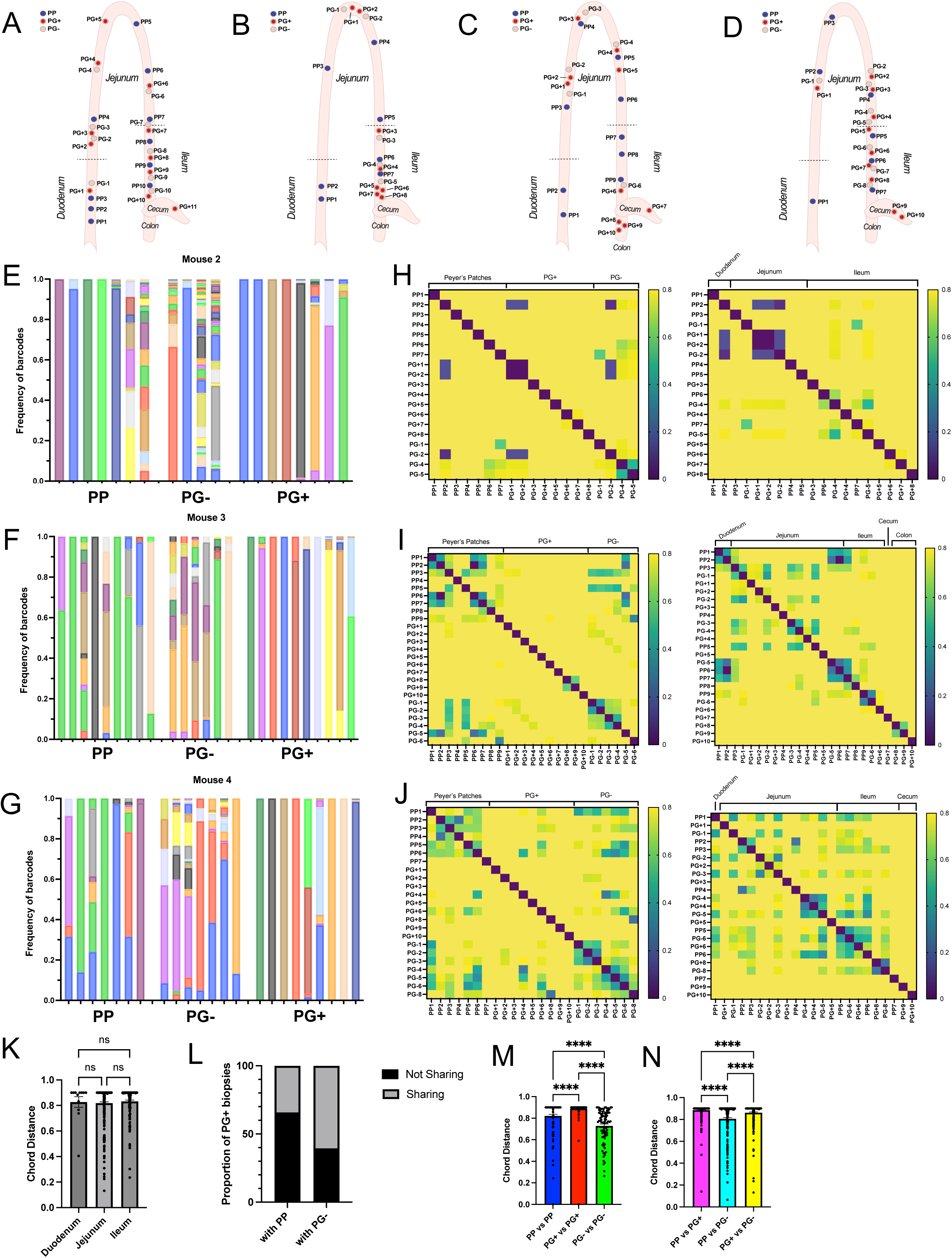
Intestinal biopsy maps, analysis of barcodes in mouse intestine, and pooled comparisons of chord distance. (A-D) Map showing a representative layout of individually harvested and processed tissue biopsies from the small intestine, cecum, and colon of mice 1-4 on day 5 post-infection. (E-G) Frequency of barcodes per biopsy for mice 2-4 (mouse 1 is depicted in Fig. 1C), where each bar represents one biopsy, and each color represents one barcode. (H-J) Similarity between *Y. ptb* populations in each biopsy as assessed by chord distance (CD) for mice 2-4 (mouse 1 is depicted in Fig. 1D), where (LEFT) data is organized by biopsy type and (RIGHT) data is ordered by location along gastrointestinal tract. (K) CD between biopsies within the indicated small intestinal region. Each circle represents one comparison between biopsies within the indicated region for that mouse. (L) Proportion of PG+ biopsies that share or do not share with PP or PG- biopsies in the same mouse. Sharing was determined by presence of single PG+ founder in other biopsies or, for PG+ containing more than one founder, the most abundant barcode was used to determine sharing. (M-N) CD between the indicated biopsy types. Each circle represents one comparison between the (M) same or (N) different biopsy types as indicated, where biopsies are matched within mouse. Unless otherwise indicated, for all data bars represent mean ± SEM, (pooled for n=4 mice, one experiment collected at 5 days post-infection). Statistical significance was determined using one-way ANOVA with post-hoc Dunn’s Multiple Comparison Test, where ns = not significant and **** p<0.0001.

**Supplemental Figure 3.**
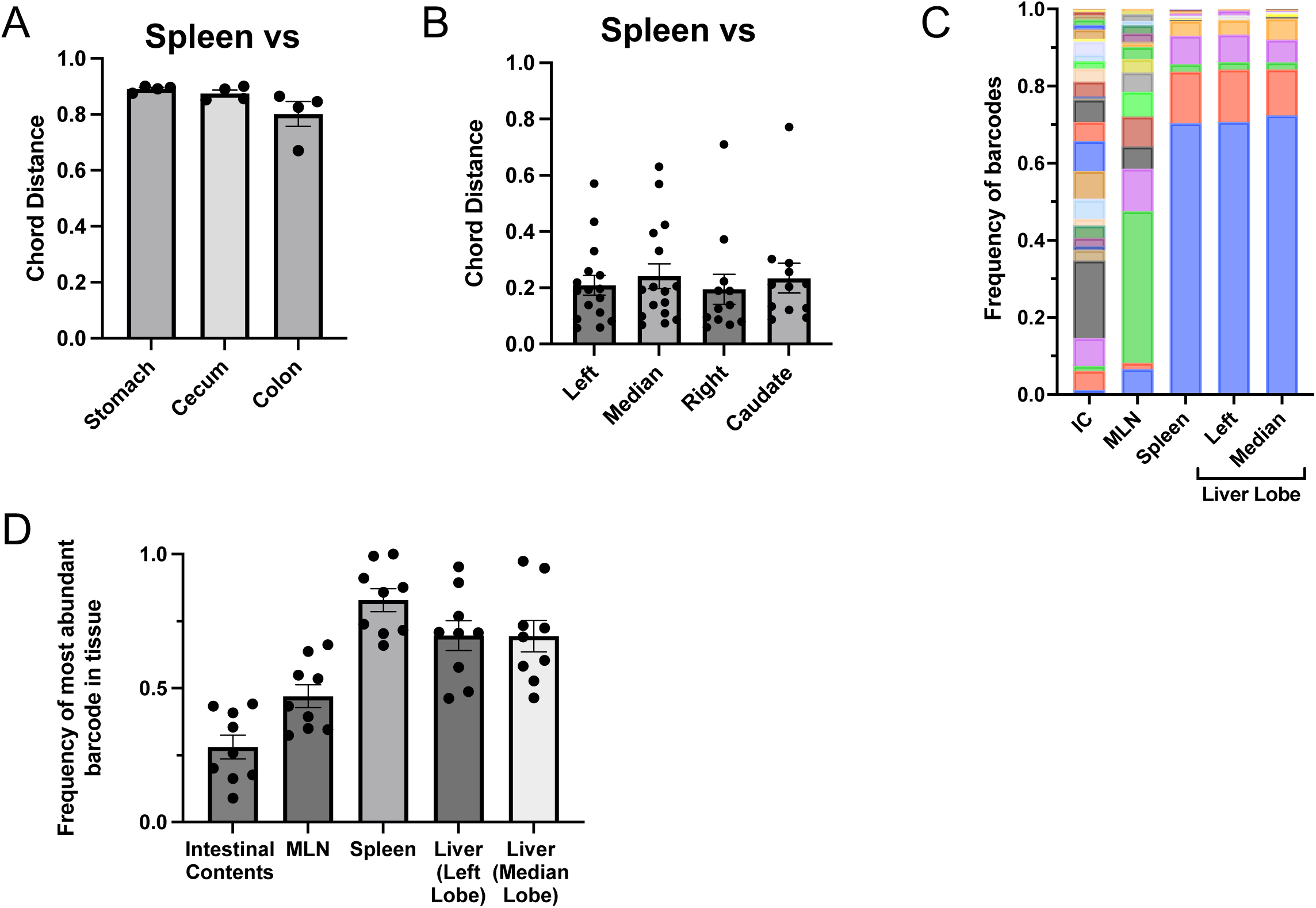
Populations of *Y. ptb* in spleen and liver are similar to each other with one highly abundant clone. (A) CD between the spleen and gastrointestinal tract organ tissues. (B) CD between the spleen and individual liver lobes. (C) Frequency of barcodes per tissue for one representative mouse. Each bar represents one sample, and each color represents one barcode. (D) Frequency of most abundant barcode per sample. All samples collected at 5 days post-infection. Data are pooled from three to five experiments, each circle represents one mouse, bars are mean ± SEM unless otherwise indicated.

**Supplemental Figure 4.**
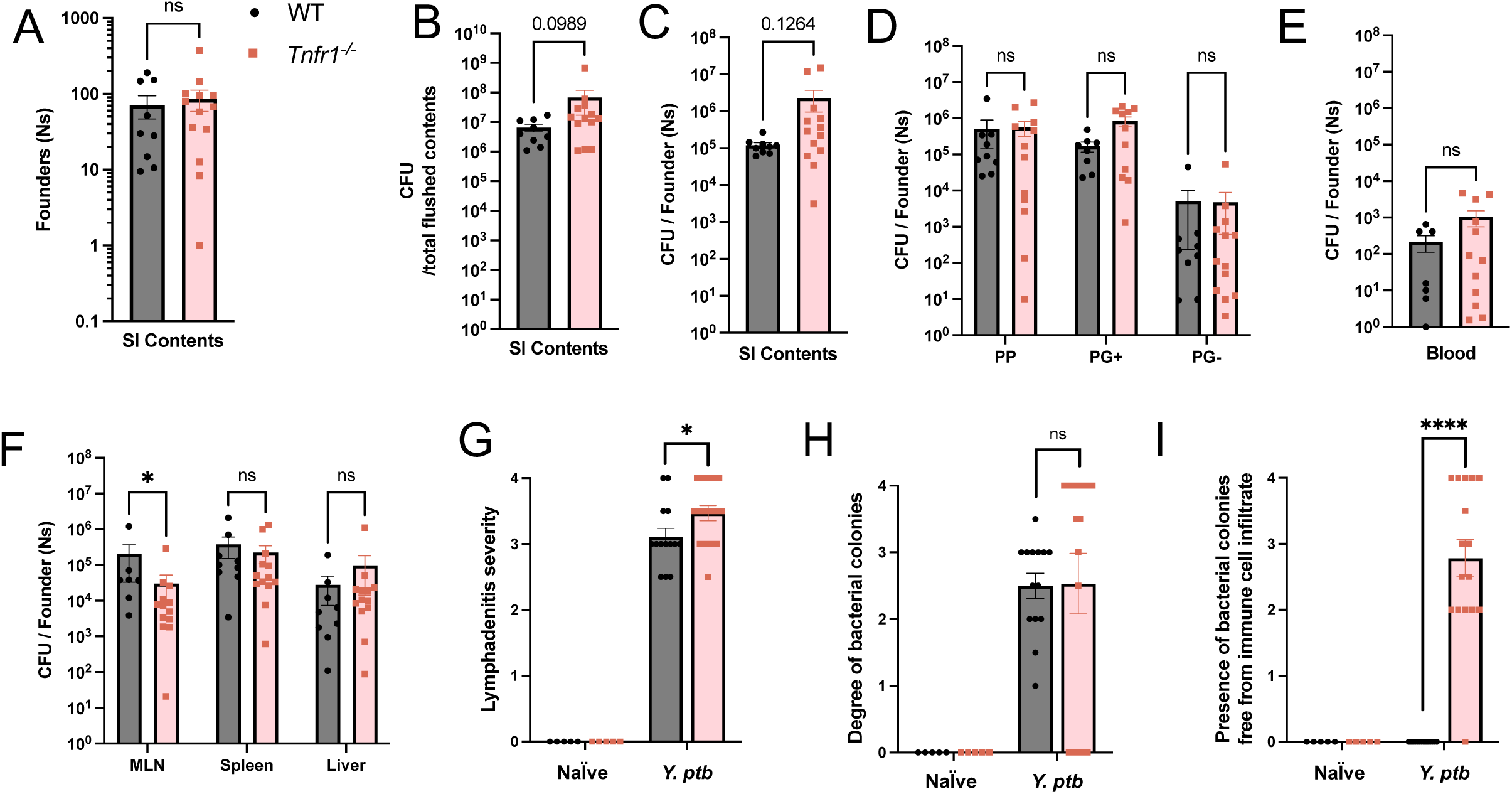
TNFR1 signaling affects bacterial colony containment in mesenteric lymph nodes but not clone expansion or lymphadenitis. (A) Bacterial burdens and (B) founding populations (Ns) in flushed small intestinal contents at day 5 post-infection. (C-F) Bacterial burden per founder, as determined by CFU/Ns, in (C) flushed small intestinal contents, (D) small intestinal PP, PG+, and PG- biopsies, (E) whole blood, and (F) indicated systemic tissues at day 5 post-infection. Liver samples show data from the median lobe as representative of all liver lobes. Pooled data from three independent experiments. (G-I) H&E-stained paraffin-embedded mesenteric lymph node sections from naïve and *Y.ptb*-infected WT and *Tnfr1^−/−^* mice at day 5 after infection were scored between 0 and 4 (minimal to extensive) for the metrics shown. Histological data are from two independent experiments. Each symbol represents one mouse, and bars are mean ± SEM. Statistical significance was determined using multiple Mann-Whitney tests and ns = not significant, * p<0.05, ** p<0.01.

